# Loss-of-function mutation in Omicron variants reduces spike protein expression and attenuates SARS-CoV-2 infection

**DOI:** 10.1101/2023.04.17.536926

**Authors:** Michelle N. Vu, R. Elias Alvarado, Dorothea R. Morris, Kumari G. Lokugamage, Yiyang Zhou, Angelica L. Morgan, Leah K. Estes, Alyssa M. McLeland, Craig Schindewolf, Jessica A. Plante, Yani P. Ahearn, William M. Meyers, Jordan T. Murray, Patricia A. Crocquet-Valdes, Scott C. Weaver, David H. Walker, William K. Russell, Andrew L. Routh, Kenneth S. Plante, Vineet Menachery

**Affiliations:** Microbiology and Immunology, University of Texas Medical Branch, Galveston, TX, United States; Institute for Translational Sciences, University of Texas Medical Branch, Galveston, TX, United States; Pediatrics, University of Texas Medical Branch, Galveston, TX, United States; Biochemistry and Molecular Biology, University of Texas Medical Branch, Galveston, TX, United States; Pathology, University of Texas Medical Branch, Galveston, TX, United States; Institute for Human Infection and Immunity, University of Texas Medical Branch, Galveston, TX, United States; World Reference Center of Emerging Viruses and Arboviruses, University of Texas Medical Branch, Galveston, TX, United States; Center for Biodefense and Emerging Infectious Disease, University of Texas Medical Branch, Galveston, TX, United States

## Abstract

SARS-CoV-2 Omicron variants emerged in 2022 with >30 novel amino acid mutations in the spike protein alone. While most studies focus on receptor binding domain changes, mutations in the C-terminus of S1 (CTS1), adjacent to the furin cleavage site, have largely been ignored. In this study, we examined three Omicron mutations in CTS1: H655Y, N679K, and P681H. Generating a SARS-CoV-2 triple mutant (YKH), we found that the mutant increased spike processing, consistent with prior reports for H655Y and P681H individually. Next, we generated a single N679K mutant, finding reduced viral replication *in vitro* and less disease *in vivo.* Mechanistically, the N679K mutant had reduced spike protein in purified virions compared to wild-type; spike protein decreases were further exacerbated in infected cell lysates. Importantly, exogenous spike expression also revealed that N679K reduced overall spike protein yield independent of infection. Although a loss-of-function mutation, transmission competition demonstrated that N679K had a replication advantage in the upper airway over wild-type SARS-CoV-2 in hamsters, potentially impacting transmissibility. Together, the data show that N679K reduces overall spike protein levels during Omicron infection, which has important implications for infection, immunity, and transmission.

## Introduction

Since its introduction, SARS-CoV-2 has continuously evolved giving rise to multiple Variants of Concern (VOCs) with diverse mutations in the spike protein ^1^ (**Extended Fig. 1A**). Present as a trimer on virions, spike is composed of S1 and S2 subunits, responsible for receptor binding and membrane fusion, respectively ^2,3^. The S1 subunit contains the N-terminal domain (NTD), receptor binding domain (RBD), and the C-terminus of the S1 subunit (CTS1), which harbors a furin cleavage site (FCS) in SARS-CoV-2. Following receptor binding, the spike is cleaved at the S1/S2 site by host proteases to expose the fusion machinery for entry. With the diverse mutations in the spike protein (**Extended Fig. 1B**), most Omicron studies have focused on the RBD and the impact on vaccine- or infection-induced immunity. However, mutations surrounding the FCS and S1/S2 cleavage site have been demonstrated to drive SARS-CoV-2 pathogenesis ^4-9^ and have been largely unstudied in the context of Omicron.

With this in mind, we set out to evaluate the role of Omicron CTS1 mutations on infection and pathogenesis. Omicron maintains three mutations adjacent to the FCS and S1/S2 cleavage site: H655Y, N679K, and P681H (**Extended Fig. 1B**). Both H655Y and P681H have previously been observed in the Gamma and Alpha variants ^10,11^; in contrast, N679K is unique to and maintained by all Omicron subvariants ^12^. To evaluate the role of these mutations, we used reverse genetics to generate SARS-CoV-2 mutants with all three CTS1 mutations (YKH) or N679K alone in the original WA1 backbone from early 2020. While YKH modestly increases viral replication and spike processing *in vitro*, N679K results in a loss-of-function mutation that attenuates viral replication *in vitro* and disease *in vivo* while skewing replication toward the upper airways through reduced spike protein expression. Given the importance of spike protein for immunity, our finding may have major implications for vaccine efficacy and breakthrough infections.

## Results

### H655Y, N679K, and P681H together increase viral replication and spike processing

While the majority of the > 30 spike mutations Omicron acquired are localized to the RBD, three are harbored in the CTS1 adjacent to the furin cleavage site – H655Y, N679K, and P681H (**Fig. 1A**). Both H655Y and P681H have been observed individually in Gamma and Alpha variants and are associated with increased spike processing. In contrast, N679K is a mutation unique to Omicron and is maintained in all subsequent Omicron subvariants despite involving a single nucleotide change (T/C to A/G) in the wobble codon position ^12^. Importantly, N679K is adjacent to an important O-linked glycosylation site at T678 ^13,14^; our group has previously shown this glycosylation is important for SARS-CoV-2 infection and protease usage ^8^.

**Figure 1.**
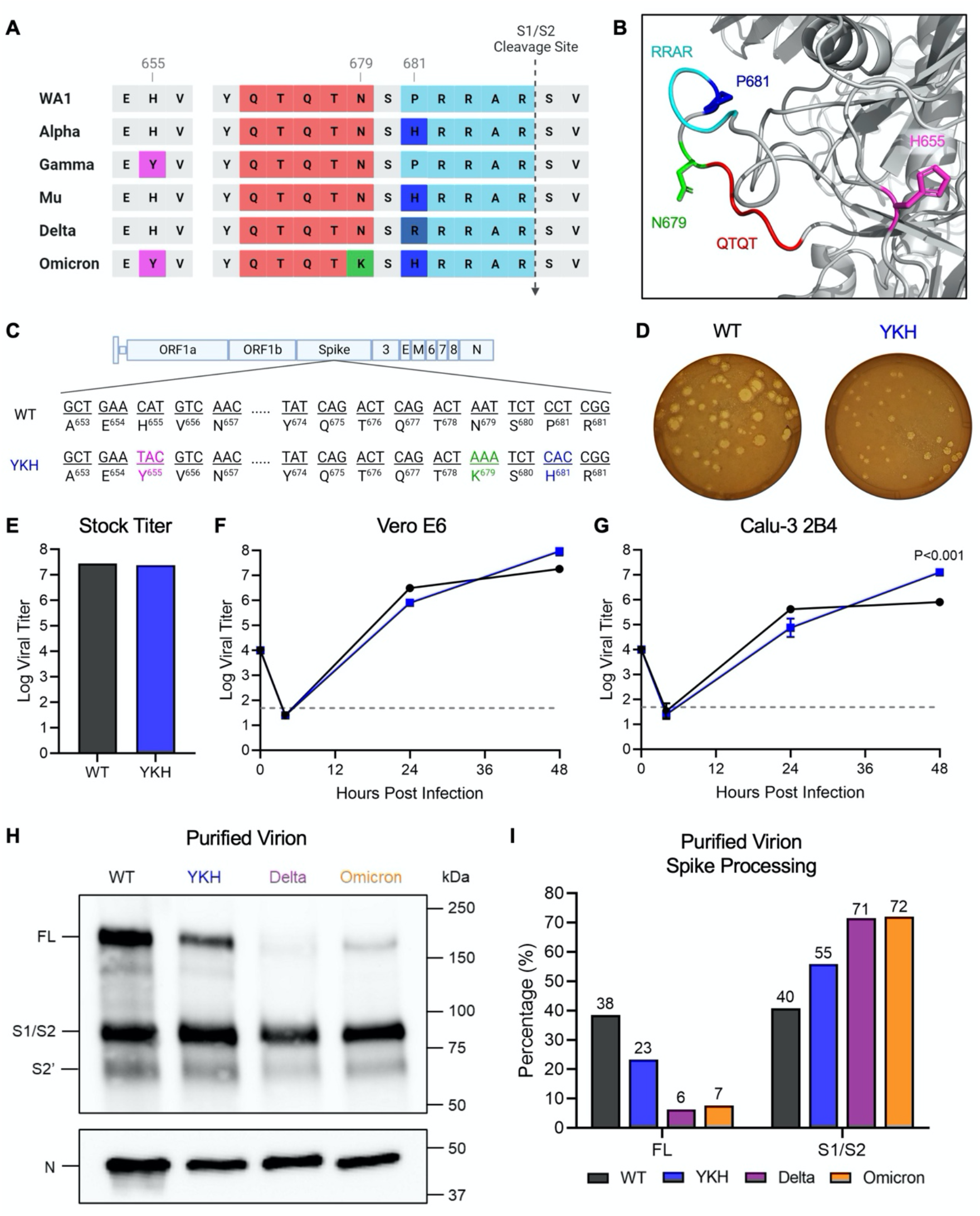
The combination of Omicron mutations H655Y, N679K, and P681H increases viral replication and spike processing. **(A)** Comparison of CTS1 region near the S1/S2 cleavage site between SARS-CoV-2 variants. **(B)** Structure of loop containing the S1/S2 cleavage site on SARS-CoV-2 spike protein. The residues that are mutated in Omicron are shown: H655 (magenta), N679 (green), and P681 (blue). The furin cleavage site RRAR (cyan) and QTQT motif (red) are also shown. **(C)** Schematic of WT and YKH SARS-CoV-2 mutant genomes. **(D)** WT and YKH SARS-CoV-2 plaques on Vero E6 cells at 2 dpi. **(E)** Viral titer from WT and YKH virus stocks representing the highest yield generated from TMPRSS2-expressing Vero E6 cells. **(F-G)** Replication kinetics of WT and YKH in Vero E6 **(F)** and Calu-3 2B4 **(G)** cells. Cells were infected at an MOI of 0.01 infectious units/cell (n=3). Data are mean ± s.d. Statistical analysis measured by two-tailed Student’s t-test. **(H)** Purified WT, YKH, Delta isolate (B.1.617.2), and Omicron (BA.1) virions from Vero E6 supernatant were probed with α-Spike and α-Nucleocapsid (N) antibodies in Western blots. Full-length spike (FL), S1/S2 cleavage product, and S2’ cleavage product are indicated. **(I)** Densitometry of FL and S1/S2 cleavage product was performed, and quantification of FL and S1/S2 cleavage product percentage of total spike is shown. Quantification was normalized to N for viral protein loading control. WT (black), YKH (blue), Delta isolate (purple), Omicron (orange). Results are representative of two experiments.

Several motifs within the CTS1 spike domain, including the furin cleavage site and the upstream QTQTN motif, are key to spike cleavage and host protease interactions, which drive SARS-CoV-2 infection and pathogenesis. All three Omicron mutations in the CTS1, H655Y, N679K, and P681H, are adjacent to or within these motifs and may impact their function (**Fig. 1A and 1B**). To evaluate this, we generated a mutant SARS-CoV-2 harboring H655Y, N679K, and P681H in the original WA1 backbone (YKH) (**Fig. 1C**) ^15,16^. Plaques produced by the YKH mutant were smaller compared to the parental WA1 (WT) (**Fig. 1D**). However, the YKH mutant did not attenuate stock titers nor replication kinetics in Vero E6 cells as compared to wild-type (WT) SARS-CoV-2 (**Fig. 1E and 1F**). Notably, while replication was slightly reduced at 24 hpi, end point titers for YKH were augmented at 48 hpi in Calu-3 2B4 cells compared to WT (**Fig. 1G**). The results suggest that the combination of the three mutations alters infection dynamics, which may offer some advantages to the Omicron variant in human respiratory cells (**Fig. 1G**). As H655Y and P681H have individually been shown to increase spike processing, we next evaluated spike processing on purified virions from YKH and WT infection. Similar to Delta and Omicron, YKH spike was more processed than WT (**Fig. 1H and 1I**). At 24 hpi, the S1/S2 cleavage ratio to full length spike ratio was ∼2.4:1 for the YKH spike (55% S1/S2 product, 23% full-length); in contrast, WT had roughly equivalent amounts of S1/S2 product and full length. Overall, the combination of H655Y, N679K, and P681H in the YKH mutant resulted in increased viral endpoint yields in human respiratory cells and contributed to Omicron’s enhanced spike processing.

### N679K mutation attenuates SARS-CoV-2 infection

The increase in spike processing found in the YKH mutant is consistent with prior work examining H655Y and P681H mutations individually; however, the contribution of N679K had yet to be evaluated. Based on its location adjacent to a key O-linked glycosylation site ^8^, we hypothesized that N679K might impact SARS-CoV-2 infection (**Fig. 2A**). To evaluate potential changes, we generated a SARS-CoV-2 mutant with only N679K in the WA1 backbone (N679K) (**Fig. 2B**). Our initial characterization found that the N679K plaque sizes were distinctly smaller at days 2 and 3 post-infection (**Fig. 2C**), and stock titers were also slightly lower than WT (**Fig. 2D**). These differences in plaque size and stock titers are consistent with observations of most Omicron strains ^17-20^. Notably, unlike the minimal differences seen in YKH replication kinetics, the N679K mutant had attenuated replication in both Vero E6 and Calu-3 2B4 cells at 24 hpi (**Fig. 2E and 2F**). Although N679K viral titer recovered by 48 hpi, the results suggest that N679K is a loss-of-function mutation in terms of replication in both cell lines.

**Figure 2.**
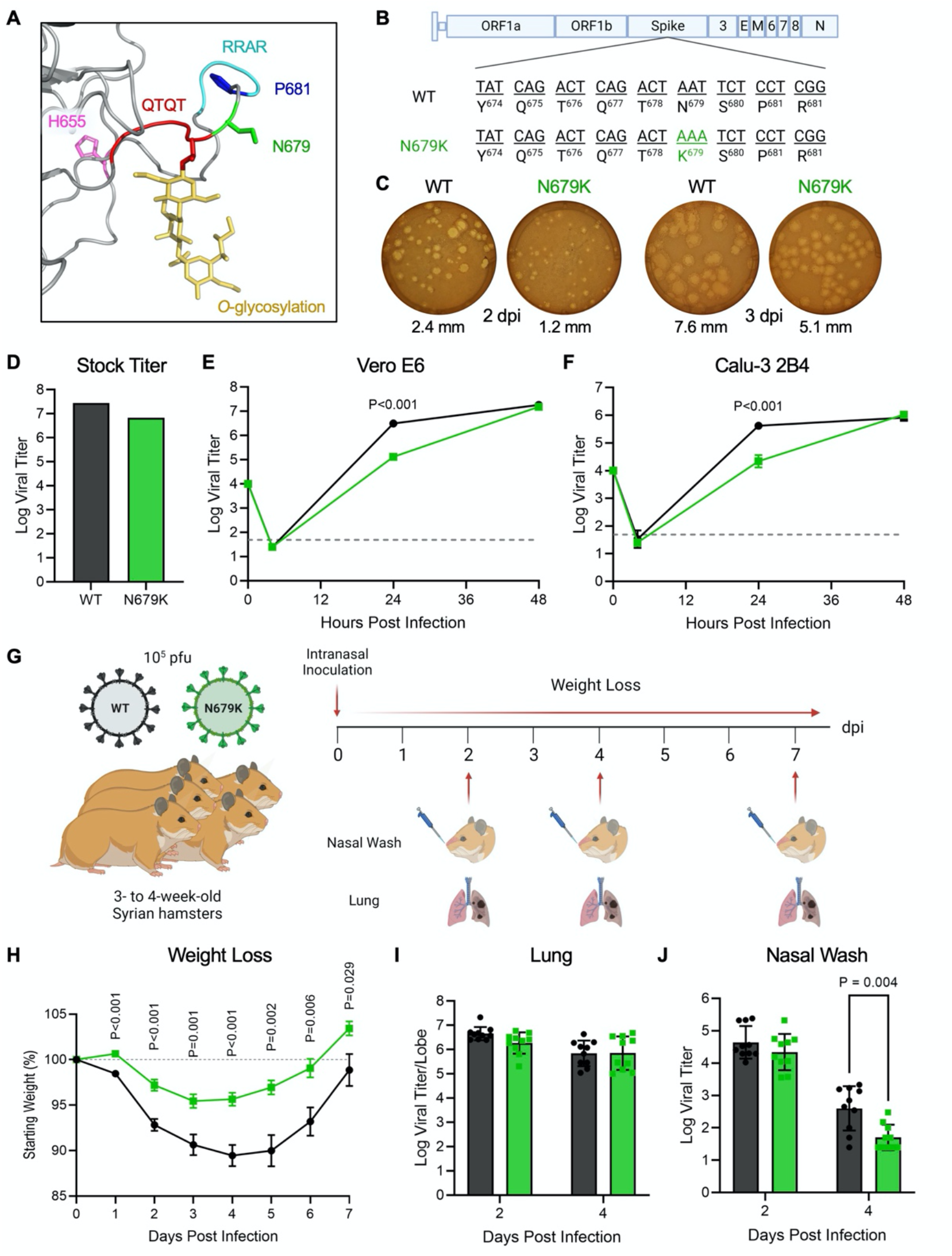
N679K attenuates SARS-CoV-2 replication and disease. **(A)** Structural modeling of O-linked glycosylation of threonine 678 (yellow) of QTQTN motif (red) and the residues mutated in Omicron – H655 (magenta), N679 (green), and P681 (blue) – with N679 adjacent to the glycosylation. The furin cleavage site RRAR is also shown (cyan). **(B)** Schematic of WT and N679K SARS-CoV-2 genomes. **(C)** WT and N679K SARS-CoV-2 plaques on Vero E6 cells at 2 (left) and 3 dpi (right). Average plaque sizes noted below. **(D)** Viral titer from WT and N679K virus stocks with the highest yield generated form TMPRSS2-expressing Vero E6 cells. **(E-F)** Replication kinetics of WT and N679K in Vero E6 **(E)** and Calu-3 2B4 **(F)** cells. Cells were infected at an MOI of 0.01 infectious units/cell (n=3). Data are mean ± s.d. Statistical analysis performed using two-tailed Student’s t-test. **(G)** Schematic of experimental design for golden Syrian hamster infection with WT (black) or N679K (green) SARS-CoV-2. Three- to four-week-old male hamsters were infected with 10^5^ pfu and monitored for weight loss over 7 days. At 2, 4, and 7 dpi, nasal washes and lungs were collected for viral titer, and lung was collected for histopathology. **(H)** Weight loss of hamsters infected with WT (black) or N679K (green) SARS-CoV-2 over 7 days. Data are mean ± s.e.m. Statistical analysis measured by two-tailed Student’s t-test. **(I-J)** Viral titers of lungs **(I)** and nasal washes **(J)** collected at 2 and 4 dpi from hamsters infected with WT (black) or N679K (green) SARS-CoV-2. Data are mean ± s.d. Statistical analysis measured by two-tailed Student’s t-test.

We next evaluated N679K *in vivo* by infecting 3-to-4-week-old golden Syrian hamsters and monitored weight loss and disease over 7 days (**Fig. 2G**). Hamsters infected with N679K displayed significantly attenuated body weight loss compared to those infected with WT (**Fig. 2H**). Despite the stark attenuation seen in weight loss, N679K viral titers in the lungs were equivalent to WT at 2 dpi and 4 dpi (**Fig. 2I**). Similarly, N679K viral titers were comparable to WT at 2 dpi in nasal washes; however, the mutant virus resulted in reduced replication at 4 dpi (**Fig. 2J**). In addition, analysis of lung histopathology showed a modest, but not significant reduction in disease of the N679K infected hamsters as compared to control (**Extended Fig. 2**). Taken together, our results indicate that N679K has a distinct loss-of-function phenotype *in vitro* and *in vivo*.

### N679K mutation results in decreased spike protein expression

We next sought to determine the mechanism driving the loss-of-function observed with the N679K mutant. Given its location adjacent to the FCS, we first evaluated N679K effects on proteolytic spike processing. Virions were purified from WT, N679K or the Omicron variant BA.1 (Omicron) and blotted for spike processing. Nearly identical to YKH, the N679K mutant had increased spike processing with a ∼2.5:1 ratio of S1/S2 cleavage product to full-length spike compared to 1:1 ratio for WT at 24 hpi (**Fig. 3A and 3B**). However, we noted distinct differences in total spike protein with N679K and Omicron compared to WT, despite similar levels of nucleocapsid protein. Densitometry analysis revealed that the total spike to nucleocapsid (S/N) ratio of N679K and Omicron virions was reduced 21% and 36%, respectively, as compared to WT (**Fig. 3C**). Overall, our results indicate that the N679K mutant and Omicron variant incorporate less spike protein into their virions.

**Figure 3.**
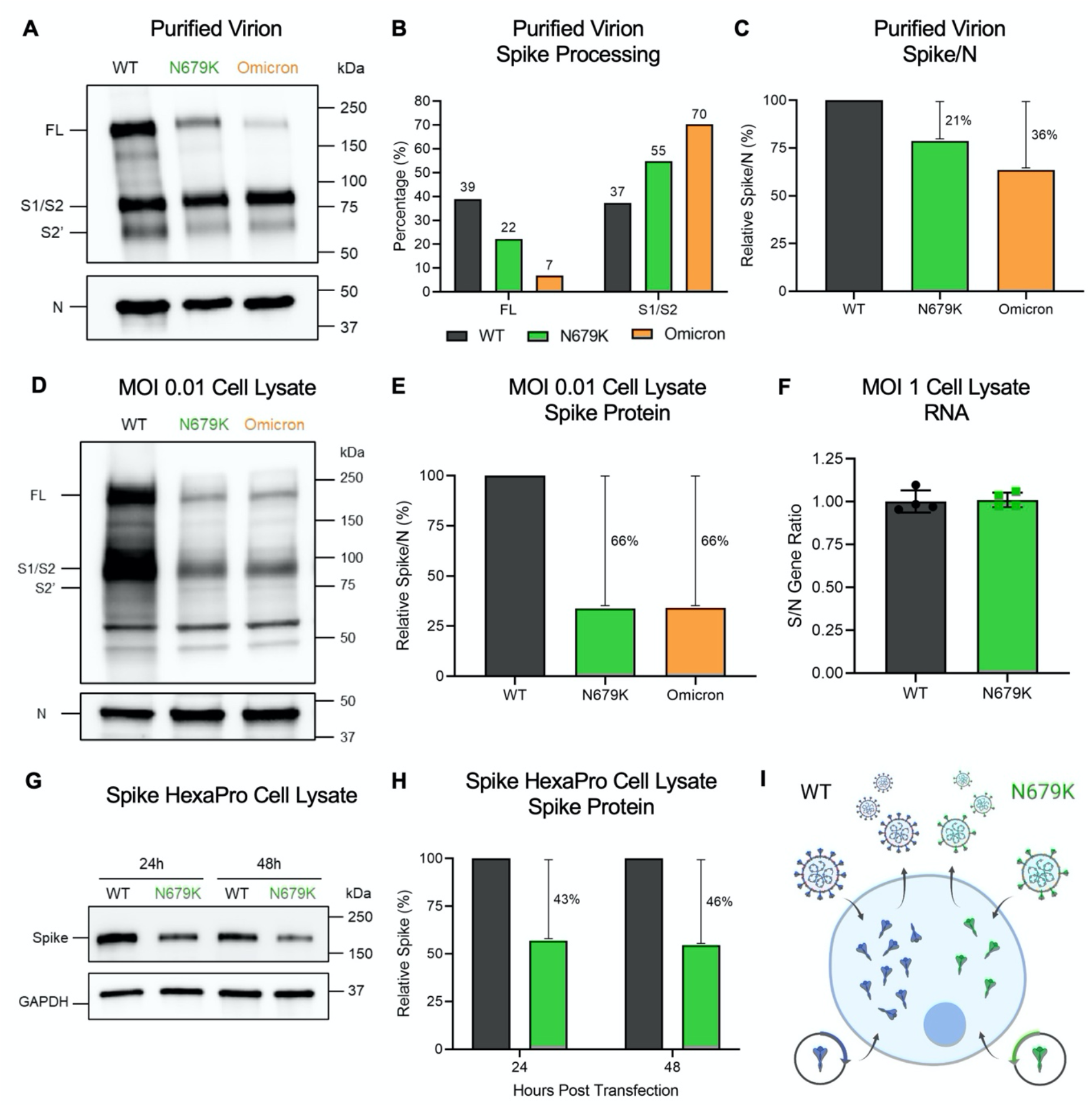
N679K results in decreased spike expression on virions and in infected cells. **(A)** Purified WT, N679K, and Omicron (BA.1) virions from Vero E6 supernatants were probed with α-Spike and α-Nucleocapsid (N) antibodies in Western blots. Full-length spike (FL), S1/S2 cleavage product, and S2’ cleavage product are indicated. **(B)** Densitometry of spike processing from purified virions applied to Western blots in **(A)** was performed, and quantification of FL and S1/S2 cleavage product percentage of total spike is shown. Quantification was normalized to N as viral protein loading control. WT (black), N679K (green), Omicron (orange). Results are representative of two experiments. **(C)** Densitometry of spike expression from purified virion Western blots in **(A)** was performed, and quantification of total spike protein to nucleocapsid ratio is shown. Spike/N ratio is relative to WT. WT (black), N679K (green), Omicron (orange). Results are representative of two experiments. **(D)** Vero E6 cells were infected with WT, N679K, or Omicron at an MOI of 0.01 infectious units/cell. Cell lysate was collected at 24 hpi and probed with α-Spike and α-Nucleocapsid (N) antibodies in Western blots. Full-length spike (FL), S1/S2 cleavage product, and S2’ cleavage product are indicated. **(E)** Densitometry of spike expression from infected cell lysate Western blots in **(D)** was performed, and quantification of total spike protein to nucleocapsid ratio is shown. Spike/N ratio is relative to WT. WT (black), N679K (green), Omicron (orange). Results are representative of three biological replicates. **(F)** Vero E6 cells were infected with WT or N679K at an MOI of 1 infectious units/cell. Cell lysate was collected at 8 hpi in Trizol to extract RNA. RNA transcripts for spike, nucleocapsid and 18S were measured using RT-qpCR. The ratios of ΔΔCt spike to ΔΔCt nucleocapsid are shown. Data are mean ± s.d. Statistical analysis measured by two-tailed Student’s t-test. **(G)** Vero E6 cells were transfected with Spike HexaPro WT and N679K and cell lysate was collected at 8, 24, and 48 hpt. Lysates were probed with α-Spike and α-GAPDH antibodies in Western blots. **(H)** Densitometry of spike expression from transfected cell lysates by Western blot in **(G)** was performed, and quantification of relative total spike protein is shown. Spike protein levels were normalized to GAPDH and are relative to WT. WT (black), N679K (green). Results are representative of three biological replicates. **(I)** While WT virus and exogenous spike plasmid produces abundant spike protein, the N679K mutation results in less spike protein expression in virions, intracellularly by infection and transfection of exogenous spike plasmid.

We then sought to determine if changes in the virion spike were due to changes to total protein expression in the cell or spike incorporation into the particle. To examine spike protein expression, we measured total spike relative to nucleocapsid from infected Vero E6 cell lysates 24 hpi (**Fig. 3D and 3E**). N679K resulted in a S/N ratio 66% less than WT, displaying an even further decrease in spike protein compared to the reduction in purified virions. Additionally, a similar decrease in S/N ratio was observed in Omicron, indicating that the phenotype is maintained in the context of all the Omicron mutations (**Fig. 3D and 3E**). Importantly, the RNA transcript ratio for both spike and N following infection of WT and N679K were nearly identical indicating no deficits in RNA expression of spike in the mutant (**Fig. 3F**). Together, the results indicate that the N679K mutation reduces the Omicron spike protein levels compared to WT following infection.

Having established reduced spike protein in the context of N679K, we next wanted to determine if this reduction only occurs in the context of virus infection or is inherent to the protein. Therefore, we introduced the mutation into the Spike HexaPro plasmid to exogenously express spike protein and separate N679K driven changes from other aspects of viral infection ^21^. Vero E6 cells were transfected with the WT or N679K mutant spike HexaPro and harvested at 24 and 48 hours post transfection (hpt). Similar to what was observed in viral infection, N679K spike was reduced 43% at 24 hpt and 46% at 48 hpt (**Fig. 3G and 3H**). Overall, the results across virions, cell lysates, and overexpression systems demonstrate that the reduction in spike protein is governed by the N679K mutation in a manner independent of viral infection (**Fig. 3I**).

### N679K mutation results in preference for the upper airways

Recognizing that decreased spike expression impacts virus infection, we next evaluated the role of N679K on SARS-CoV-2 transmissibility. Using transmission competition, donor hamsters were infected with a 1:1 ratio of WT:N679K SARS-CoV-2 at a total of 10^5^ pfu **(Fig. 4A)**. At 24 hpi, donors were paired with naïve recipients, cohoused for 8 hrs, separated, and donors nasal washed. Nasal wash, trachea, and lung were collected to measure viral RNA populations at 2 and 4 days post infection (dpi) for donors and post contact (dpc) for recipients. Surprisingly, while both viruses transmitted, WT and N679K demonstrated distinct replication sites along the respiratory tract **(Fig. 4B)**. N679K dominated the nasal washes and upper airways while WT primarily seeded the lungs and lower airways. The trachea serves as a midpoint, with no clear delineation between the viruses **(Fig. 4B)**. Having observed this gradation, we returned to the prior hamster study and examined antigen staining of the lung **(Fig. 4C)**. While no significant differences in total antigen were noted, the localization of viral antigen in the N679K infection was distinct and concentrated in the large airways. In contrast, WT was more uniformly distributed in the parenchyma and airways **(Fig. 4D)**. Overall, the results indicate that the N679K mutation shifts viral replication towards airway replication.

**Figure 4.**
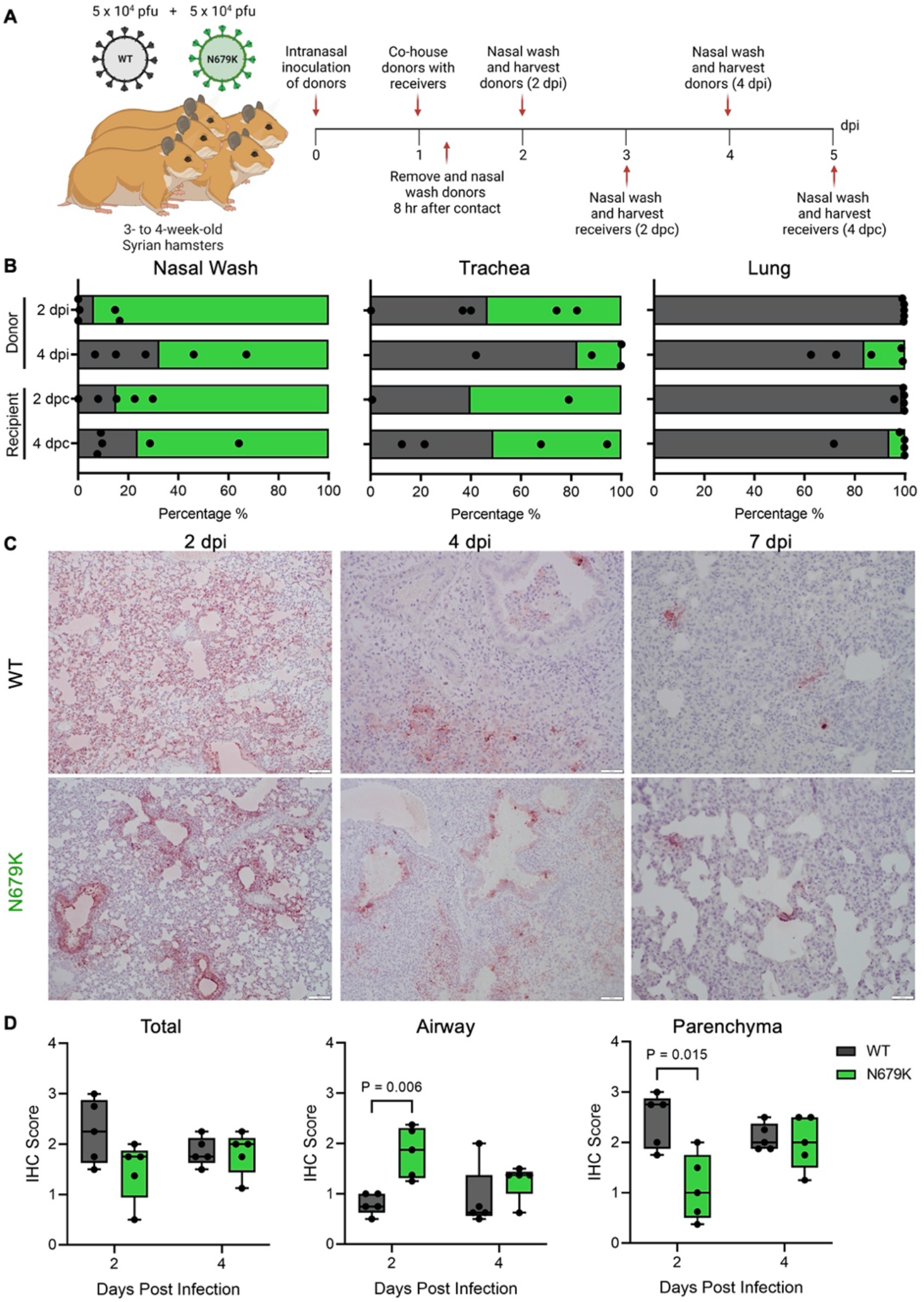
N679K results in preference for upper airways. **(A)** Schematic of experimental design of transmission competition in golden Syrian hamsters. Donor three- to four-week-old male hamsters were intranasally infected with 10^5^ pfu of WT:N679K SARS-CoV-2 in a 1:1 ratio and housed singly. Donors were paired with recipients 24 hpi and cohoused for 8 hrs before separating and nasal washing donors. Nasal washes, tracheas, and lungs were collected at 2 and 4 days post infection for donors (dpi) and post contact for recipients (dpc). **(B)** Next generation sequencing was performed on extracted RNA to measure the percentage of WT (black) and N679K (green) present in nasal wash (left), trachea (middle), and lung (right) of donors (top) and recipients (bottom). **(C)** Immunohistochemistry of left lung lobes at 2, 4 and 7 dpi staining for nucleocapsid. Hamsters were singly infected with 10^5^ pfu of either WT or N679K SARS-CoV-2. **(D)** Immunohistochemistry staining of left lung lobes form hamsters infected with WT (black) or N679K (green) SARS-CoV-2 were scored by total section (left), airway (middle), or parenchyma (right) staining. Data are mean showing minimum and maximum (n=5). Statistical analysis measured by two-tailed Student’s t-test.

## Discussion

Most Omicron studies have focused on determining the impact that the RBD mutations have on immune escape, largely overlooking mutations in other spike domains like the CTS1. Harboring the FCS and S1/S2 cleavage site, the CTS1 has been demonstrated as a hotspot for both attenuating and augmenting mutations ^4-9^. Focusing on Omicron’s three CTS1 mutations –H655Y, N679K, and P681H, we generated infectious clones with all three (YKH) or N679K alone in the SARS-CoV-2 WA1 background. The combination of YKH produced a modest increase in endpoint titers after infection of human respiratory cells, and augmented spike processing, consistent with prior studies that tested the effects of H655Y and P681H individually ^10,11^. However, the N679K mutant reduced viral replication *in vitro* and weight loss *in vivo*. Mechanistic studies determined that both N679K and Omicron have reduced spike protein incorporated into their virions, less spike protein in infected cell lysates, and inferior production using exogenous spike protein expression systems. Our results argue that reduced spike protein in the context the N679K mutation attenuates Omicron strains and may have implications for SARS-CoV-2 immunity by reducing spike antigen thus shifting immune recognition. Additionally, while the N679K mutation attenuated the virus *in vivo*, our studies indicate a shift toward the upper airways replication. Overall, the data argue that N679K acts as a loss-of-function mutation that has a significant impact on SARS-CoV-2 Omicron infection, pathogenesis, and transmission dynamics.

N679K is likely attenuated because of its decrease in spike protein production. Starting with ∼20-30% less spike in its virions, one possibility was a change in spike incorporation. However, an even greater decrease (66%) in spike protein was present in infected Vero E6 cell lysates, indicated that overall spike protein levels were affected. In addition, we found no change in the ratio of spike message relative to N transcript, suggesting the N679K mutation impacts the protein itself. To confirm that the reduction in spike was not a product of virus infection or host immune interactions, we exogenously expressed the spike protein to demonstrate that the N679K spike protein itself was less stable than the WT control. One possible mechanism is that the asparagine-to-lysine change introduces a ubiquitination site that could lead to spike degradation. Another possible mechanism is that the N679K mutation itself may destabilize the protein structurally. Additionally, the N679K substitution adds another basic amino acid to the stretch including the furin cleavage site; the positively charged lysine extends the polybasic cleavage motif and may facilitate cleavage by additional host proteases ^22^. Overall, while the exact mechanism is unclear, the N679K mutation results in a less stable spike protein that impacts infection and pathogenesis of SARS-CoV-2.

Surprisingly, N679K is uniformly found in 100% of Omicron sequences in GSAID, despite being a single nucleotide change in the wobble position ^12^. Though attenuated *in vitro*, N679K does replicate to similar titers as WT in hamster lungs and at day 2 in nasal washes. Notably, N679K outcompetes WT in the upper airways when in direct transmission competition. These results suggest no deficits in transmission and may augment spread as virus replication in the upper airway is more likely to seed new infections. These results also potentially explain why N679K is maintained despite clear attenuation of SARS-CoV-2 infection. Notably, addition of H655Y and P681H in the YKH mutant rescues replication in Calu-3 cells, suggesting that other Omicron mutations may compensate for N679K. However, it is unclear if reverting N679K in the Omicron strains would result in a gain in terms of *in vitro* replication or *in vivo* pathogenesis. While N679K in SARS-CoV-2 WA1 produces a clear loss-of-function, the constellation of spike mutations and epistatic interactions may mitigate the deficit in Omicron strains. Importantly, the complete conservation of N679K in Omicron also implies some fitness advantage ^23-28^. From our data, the shift toward upper airway replication by N679K may explain how it is maintained despite lower overall spike protein expression.

In addition to impacting primary infection, the reduction in spike protein may have important implications for SARS-CoV-2 and human immunity. Compared to WT, the N679K mutant produces less spike protein upon infection and can potentially skew the ratio of antibodies targeting spike and nucleocapsid. Prior work with SARS-CoV had shown that an altered spike/nucleocapsid antibody ratio contributed to vaccine failure in aged mice ^29^. Therefore, infection with Omicron could increase N targeting antibodies at the expense of spike antibodies. The result would be less protective neutralizing antibody, which may facilitate more breakthrough infections. Furthermore, SARS-CoV-2 vaccines based on the Omicron spike may produce less spike protein due to N679K mutation. In the context of the mRNA bivalent vaccines, the N679K mutation may alter the 1:1 ratio of WT to Omicron spike protein; N679K may bias immune responses towards WT spike protein instead of equally between both spike proteins. In addition, the total amount of spike protein produced may be less than previous vaccines formulations, thus diminishing the overall antibody response. These factors potentially contribute to the less than expected increase in immunity against Omicron strains despite the new bivalent vaccine formulations. Moving forward, reverting K679 back to N679 in vaccine may improve spike protein yields and subsequently improve vaccine response to the Omicron variants.

Together, our results demonstrate that Omicron N679K is a loss-of-function mutation consistently maintained in subvariants. Mechanistically, the N679K mutation attenuates the virus *in vitro* and *in vivo* by increasing spike degradation. While the N679K mutation is attenuating in isolation, other Omicron mutations like H655Y and P681H may compensate for the N679K loss of function by amplifying spike processing and infection. However, the decreased spike protein expression by N679K may have implications for immunity induced by infection and vaccines. In addition, while N679K attenuated viral pathogenesis, the shift to the upper airway replication may have enhanced transmissibility and contribute to Omicron emergence. Overall, the data highlight that the Omicron CTS1 mutations have a significant impact on SARS-CoV-2 infection and are worthy of continued study and surveillance.

## Acknowledgments

Research was supported by grants from NIAID of the NIH to (R01-AI153602 and R21-AI145400 to VDM; R24-AI120942 (WRCEVA) to SCW). ALR was supported by a UTMB Institute for Human Infection and Immunity grant and the Sealy and Smith Foundation. Research was also supported by STARs Award provided by the University of Texas System to VDM and Data Acquisition award provided by the Institute for Human Infections and Immunity at UTMB to MNV. Trainee funding provided by NIAID of the NIH to MNV (T32-AI060549).

Mass spectrometry experiments and analysis was provided by the Mass Spectrometry Facility at the University of Texas Medical Branch (https://www.utmb.edu/MSF). Figures were created with BioRender.com

## Competing Interest Statement

VDM has filed a patent on the reverse genetic system and reporter SARS-CoV-2. MNV and VDM have filed a provisional patent on a stabilized SARS-CoV-2 spike protein. Other authors declare no competing interests.

## Author contributions

Conceptualization: MNV, VDM

Formal analysis: MNV, VDM

Funding acquisition: MNV, ALR, SCW, VDM

Investigation: MNV, REA, DRM, KL, CS, JAP, ALM, LKE, AMM, YPA, WMM, PAVC, JM, DHW, KP, ALR

Methodology: MNV, KSP, VDM

Project Administration: MNV, VDM

Supervision: MNV, SCW, DHW, KSP, VDM

Visualization: MNV, DHW, VDM

Writing – original draft: MNV, VDM

Writing – review and editing: MNV, REA, CS, DHW, VDM, SCW

## Figure Legends

**Extended Figure 1.**
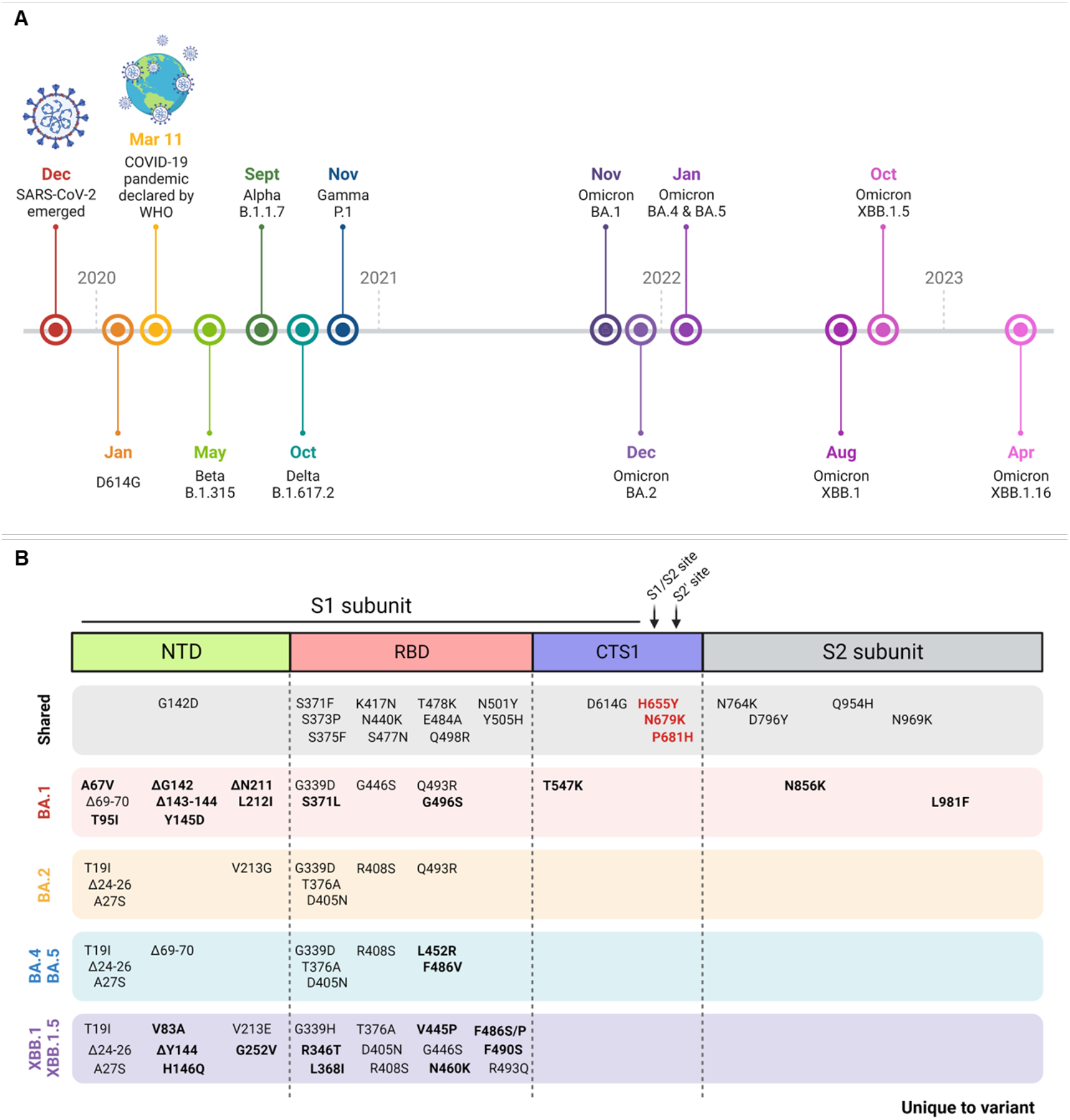
Emergence of Omicron subvariants. **(A)** Timeline of SARS-CoV-2 variants emergence by earliest documented case reported by the WHO. **(B)** Spike mutations across Omicron subvariants with shared mutations across all subvariants (gray box) and mutations unique to the specific variant (bolded) indicated. Mutations key to this study indicated in bold red.

**Extended Figure 2.**
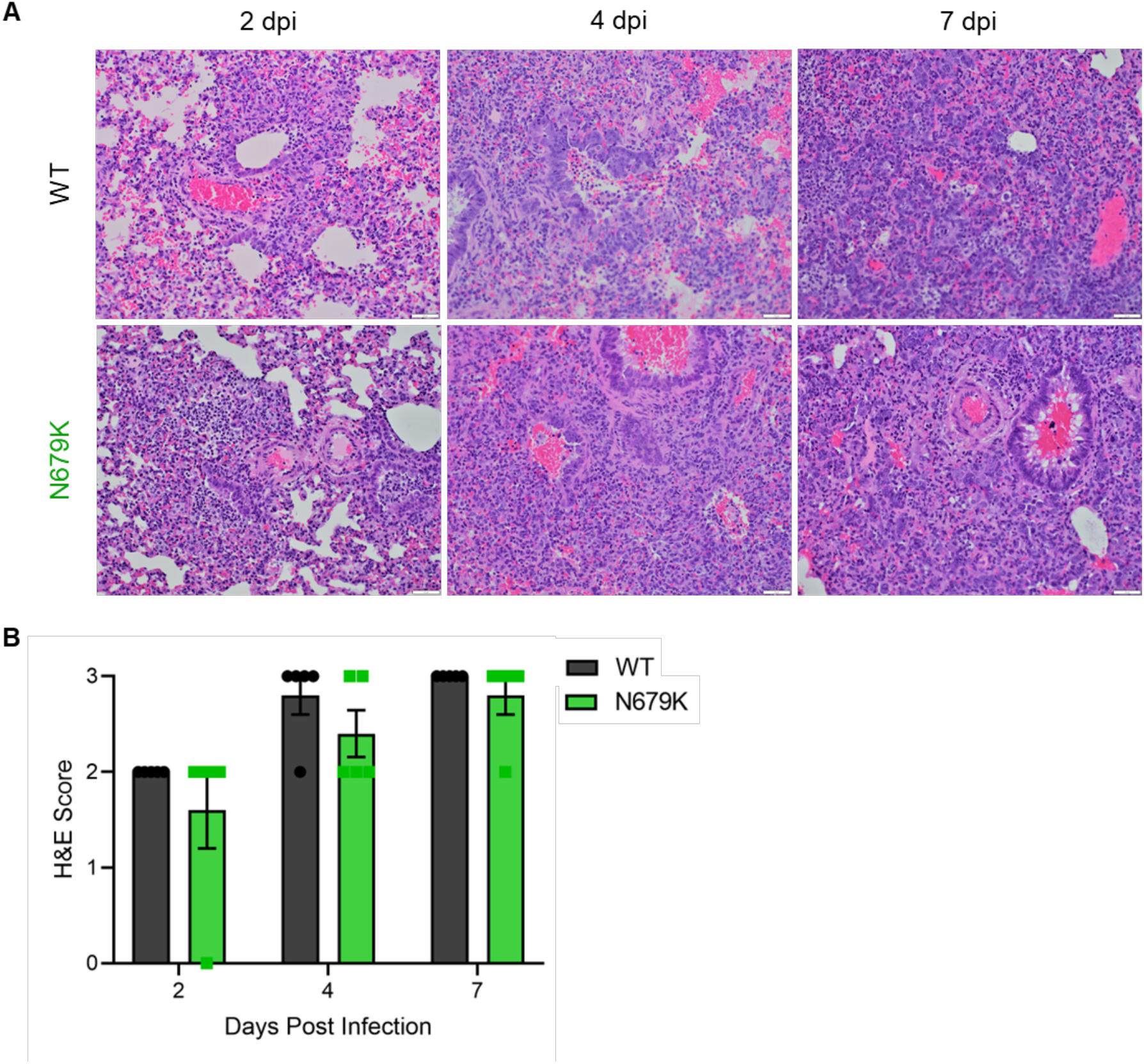
Histopathology of hamsters infected with WT or N679K SARS-CoV-2. **(A)** H&E staining of left lung of hamsters infected with 10^5^ pfu of WT (top) or N679K (bottom) SARS-CoV-2 at 2 (left), 4 (middle), and 7 (right) dpi. Lungs for both WT and N679K show bronchiolitis and interstitial pneumonia at 2 dpi that become more severe at 4 and 7 dpi. **(B)** H&E staining of left lung of hamsters infected with 10^5^ pfu of WT (black) or N679K (green) were scored for histopathological analysis.

## Methods

### Cell Culture

Vero E6 cells were grown in high glucose DMEM (Gibco #11965092) with 10% fetal bovine serum and 1x antibiotic-antimycotic. TMPRSS2-expressing Vero E6 cells were grown in low glucose DMEM (Gibco #11885084) with sodium pyruvate, 10% FBS, and 1 mg/mL Geneticin^TM^ (Invitrogen #10131027). Calu-3 2B4 cells were grown in high glucose DMEM (Gibco #11965092) with 10% defined fetal bovine serum, 1 mM sodium pyruvate, and 1x antibiotic-antimycotic.

### Viruses

The SARS-CoV-2 infectious clones were based on the USA-WA1/2020 sequence provided by the World Reference Center of Emerging Viruses and Arboviruses and the USA Centers for Disease Control and Prevention ^30^. Mutant viruses (YKH and N679K) were generated with restriction enzyme-based cloning using gBlocks encoding the mutations (Integrated DNA Technologies) and our reverse genetics system as previously described ^15,16^. Virus stock was generated in TMPRSS2-exrpressing Vero E6 cells to prevent mutations from occurring at the FCS. Viral RNA was extracted from virus stock and cDNA was generated to verify mutations by Sanger sequencing.

Delta isolate (B.1.617.2) was obtained from the World Reference Center of Emerging Viruses and Arboviruses. Infectious clone of Omicron (BA.1) was obtained from Dr. Pei Yong Shi and Dr. Xuping Xie.

### *In vitro* Infection

Vira infections in Vero E6 and Calu-3 2B4 were carried out as previously described ^8^. Briefly, growth media was removed, and cells were infected with WT or mutant SARS-CoV-2 at an MOI of 0.01 for 45 min at 37°C with 5% CO_2_. After absorption, cells were washed three times with PBS and fresh complete growth media was added. Three or more biological replicates were collected at each time point and each experiment was performed at least twice. Samples were titrated with plaque assay or focus forming assays.

### Plaque Assay

Vero E6 cells were seeded in 6-well plates and grown to 80-100% confluency in complete growth media. Ten-fold serial dilutions in PBS were performed on virus samples. Growth media was removed from cells and 200 µl of inoculum was added to monolayers. Cells were incubated for 45 min at 37°C with 5% CO_2_. After absorption, 0.8% agarose overlay was added, and cells were incubated at 37°C with 5% CO_2_ for 2 days. Plaques were visualized with neutral red stain. Average plaque size was determined using ImageJ.

### Focus Forming Assay

Focus forming assays (FFAs) were performed as previously described ^31^. Briefly, Vero E6 cells were seeded in 96-well plates to be 100% confluent. Samples were 10-fold serially diluted in serum-free media and 20 µl was to infect cells. Cells were incubated for 45 min at 37°C with 5% CO_2_ before 100 µl of 0.85% methylcellulose overlay was added. Cells were incubated for 24 h 45 min at 37°C with 5% CO_2_. After incubation, overlay was removed, and cells were washed three times with PBS before fixed and virus inactivated by 10% formalin for 30 min at room temperature. Cells were then permeabilized and blocked with 0.1% saponin/0.1% BSA in PBS before incubated with α-SARS-CoV-2 Nucleocapsid primary antibody (Cell Signaling Technology) at 1:1000 in permeabilization/blocking buffer overnight at 4°C. Cells are then washed three times with PBS before incubated with Alexa Fluor^TM^ 555-conjugated α-mouse secondary antibody (Invitrogen #A28180) at 1:2000 in permeabilization/blocking buffer for 1 h at room temperature. Cells were washed three times with PBS. Fluorescent foci images were captured using a Cytation 7 cell imaging multi-mode reader (BioTek), and foci were counted manually.

### Hamster Infection

Three- to four-week-old male golden Syrian hamsters (HsdHan:AURA strain) were purchased from Envigo. All studies were conducted under a protocol approved by the UTMB Institutional Animal Care and Use Committee and complied with USDA guidelines in a laboratory accredited by the Association for Assessment and Accreditation of Laboratory Animal Care. Procedures involving infectious SARS-CoV-2 were performed in the Galveston National Laboratory ABSL3 facility. Hamsters were intranasally infected with 10^5^ pfu of WT or N679K SARS-CoV-2 in 100 µl. Infected hamsters were weighed and monitored for illness over 7 days. Hamsters were anesthetized with isoflurane and nasal washes were collected with 400 µl of PBS on endpoint days (2, 4, and 7 dpi). Hamsters were euthanized by CO_2_ for organ collection. Nasal wash and lung were collected to measure viral titer and RNA. Left lungs were collected for histopathology.

### Transmission Competition

Three- to four-week-old male golden Syrian hamsters (HsdHan:AURA strain) were purchased from Envigo. Ten donor hamsters were intranasally infected with a 1:1 ratio of WT:N679K SARS-CoV-2 totaling 10^5^ pfu in 100 µl and were subsequently singly housed. After 24 hrs post infection, individual donor hamsters were cohoused with a recipient hamster for 8 hrs for contact transmission. Following 8 hrs, hamster pairs were separated and housed singly, and nasal washes were collected from donors. At 2 and 4 days post infection for donors and post contact for recipients, hamsters were nasal washed with 400 µl of PBS and euthanized for trachea and lung collection. Nasal washes, tracheas, and lungs were processed in TRIzol and RNA was extracted to perform next generation sequencing.

### Virion Purification

Vero E6 cells were grown in T175 flasks to be 100% confluent at time of infection. Cells were infected with 50 µl of virus stock in PBS for 45 min at 37°C with 5% CO_2_, and growth media with 5% FBS was added after absorption. Supernatant was harvested at 24 hpi and clarified by low-speed centrifugation. Virions were purified from supernatant by ultracentrifugation through a 20% sucrose cushion at 26,000 rpm for 3 hrs using a Beckman SW28 rotor. Pellets were resuspended with 2x Laemmli buffer to obtain protein samples for Western blot.

### Western Blot

Protein levels were determined by SDS–PAGE followed by western blot analysis as previously described ^8^. In brief, sucrose-purified SARS-CoV-2 virions were inactivated by resuspending in 2x Laemmli buffer and boiling. SDS-PAGE gels were run with equal volumes of samples on Mini-PROTEAN TGX gels (Bio-Rad #4561094) followed by transfer onto PVDF membrane. Membranes were incubated with α-SARS-CoV S primary antibody (Novus Biologicals #NB100-56578) at 1:1000 dilution in 5% BSA in TBST to measure spike protein processing and expression. For loading control, α-SARS Nucleocapsid primary antibody (Novus Biologicals #NB100-56576) at 1:1000 in 5% BSA in TBST was used for viral loading control and α-GAPDH primary antibody (Invitrogen #AM4300) at 1:1000 in 5% BSA in TBST for cellular loading control. Primary antibody incubation was followed by HRP-conjugated α-rabbit secondary antibody (Cell Signaling Technology #7074) or HRP-conjugated α-mouse secondary antibody (Cell Signaling Technology #7076) at 1:3000 in 5% milk in TBST. Chemiluminescence signal was developed using Clarity Western ECL substrate (Bio-Rad #1705060) or Clarity Max Western ECL substrate (Bio-Rad #1705062) and imaged with a ChemiDoc MP System (Bio-Rad). Densitometry analysis was performed using ImageLab 6.0.1 (Bio-Rad).

### RT-qPCR

Vero E6 cells were infected with an MOI of 1 as detailed above in *in vitro* infection. Cell lysate was collected at 8 hpi in TRIzol. RNA was extracted from TRIzol samples using Direct-zol RNA Miniprep Plus kit (Zymo #R2072) to be used in two-step RT-qPCR. cDNA was reverse transcribed from 1 µg of total RNA using LunaScript RT Supermix kit (NEB #E3010) according to manufacturer’s instructions. RT-qPCR was performed using Luna Universal qPCR Master mix (NEB #M3003) according to manufacturer’s instructions. RT-qPCR cycle was performed as follows: 95°C for 60 s (1 cycle), 95°C for 15 s and 51°C for 30 s then plate read (40 cycles), and melt curve from 65°C to 95°C for 5 s. For spike and nucleocapsid transcripts, a forward primer binding upstream of the transcription regulatory sequence (TRS) leader region (ACCAACCAACTTTCGATCTCT) was used with reverse primers for spike (TGCAGGGGGTAATTGAGTTCT) and nucleocapsid (CCCACTGCGTTCTCCATTCT). The 18S ribosomal RNA primers were forward (CCGGTACAGTGAAACTGCGAATG) and reverse ((GTTATCCAAGTAGGAGAGGAGCGAG). RNA transcript levels for spike and nucleocapsid were determined by ΔΔCt method with 18S as the internal control. Ratios of ΔΔCt spike over ΔΔCt nucleocapsid was reported for each sample.

### Spike HexaPro Cloning and Transfection

SARS-CoV-2 S HexaPro was a gift from Jason McLellan (Addgene plasmid #154754) ^21^. The N679K mutation was cloned into spike HexaPro using a gBlock encoding the mutation (Integrated DNA Technologies) and restriction enzyme-based cloning. Sequences were verified by Sanger sequencing.

Vero E6 cells were grown in 24-well plates to be 100% confluent at time of transfection. Cells were transfected with spike HexaPro WT or N679K plasmid and Lipofectamine 2000 following manufacturer’s instructions (Invitrogen). Briefly, 100 ng of spike HexaPro plasmid and 1.5 µl of Lipofectamine 2000 were separately diluted in 50 µl Opti-MEM (Gibco #31985070) before mixing together. After 20 min of room temperature incubation, 100 µl of the transfection mixture was added to cells, and cells were incubated at 37°C with 5% CO_2_. Cell lysate was harvested with 2x Laemmli buffer at 24 and 48 hours post transfection to be analyzed by Western blot.

### Structural Modeling

Structural models previously generated were used as a base to visualize residues mutated in Omicron ^8^. Briefly, structural models were generated using SWISS-Model to generate homology models for WT and glycosylated SARS-CoV-2 spike protein on the basis of the SARS-CoV-1 trimer structure (Protein Data Bank code 6ACD). Homology models were visualized and manipulated in PyMOL (version 2.5.4) to visualize Omicron mutations.

### Next Generation Sequencing

Next generation sequencing to determine viral RNA populations was performed as previously described ^31^. Briefly, RNA samples were extracted and prepared for Tiled-ClickSeq libraries ^32^. A modified pre-RT annealing protocol was applied as previously described^31^. The final libraries comprising of 300–700 bps fragments were pooled and sequenced on an Illumina NextSeq platform with paired-end sequencing. The raw Illumina data of the Tiled-ClickSeq libraries were processed with previously established bioinformatics pipelines^32^. One modification is the introduction of ten wild cards (“N”) covering the N679K mutation in the reference genome to allow *bowtie2^33^* to align reads to wild type or variant genomes without bias. PCR duplications were removed using *UMI-tools^34^*, and the number of unique reads representing WT and N679K variants were counted thereafter.

### Histology

Left lung lobes were harvested from hamsters and fixed in 10% buffered formalin solution for at least 7 days. Fixed tissue was then embedded in paraffin, cut into 5 µM sections, and stained with hematoxylin and eosin (H&E) on a SAKURA VIP6 processor by the University of Texas Medical Branch Surgical Pathology Laboratory.

### Immunohistochemistry

Fixed and paraffin-embedded left lung lobes from hamsters were cut into 5 µM sections and mounted onto slides by the University of Texas Medical Branch Surgical Pathology Laboratory. Paraffin-embedded sections were warmed at 56°C for 10 min, deparaffinized with xylene (3x 5-min washes) and graded ethanol (3x 100% 5-min washes, 1x 95% 5-min wash), and rehydrated in distilled water. After rehydration, antigen retrieval was performed by steaming slides in antigen retrieval solution (10 mM sodium citrate, 0.05% Tween-20, pH 6) for 40 min (boil antigen retrieval solution in microwave, add slides to boiling solution, and incubate in steamer). After cooling and rinsing in distilled water, endogenous peroxidases were quenched by incubating slides in TBS with 0.3% H_2_O_2_ for 15 min followed by 2x 5-min washes in 0.05% TBST. Sections were blocked with 10% normal goat serum in BSA diluent (1% BSA in 0.05% TBST) for 30 min at room temperature. Sections were incubated with primary anti-N antibody (Sino #40143-R001) at 1:1000 in BSA diluent overnight at 4°C. Following overnight primary antibody incubation, sections were washed 3x for 5 min in TBST. Sections were incubated in secondary HRP-conjugated anti-rabbit antibody (Cell Signaling Technology #7074) at 1:200 in BSA diluent for 1 hour at room temperature. Following secondary antibody incubation, sections were washed 3x for 5 min in TBST. To visualize antigen, sections were incubated in ImmPACT NovaRED (Vector Laboratories #SK-4805) for 3 min at room temperature before rinsed with TBST to stop the reaction followed by 1x 5-min wash in distilled water. Sections were incubated in hematoxylin for 5 min at room temperature to counterstain before rinsing in water to stop the reaction. Sections were dehydrated by incubating in the previous xylene and graded ethanol baths in reverse order before mounted with coverslips.

## Notes

### Summary of Updates

The manuscript has been modified to add figures including RNA expression data to figure 3 and transmission competition and tropism data in a new figure 4.

